# Single-cell nucleic acid profiling in droplets (SNAPD) enables high-throughput analysis of heterogeneous cell populations

**DOI:** 10.1101/2020.08.21.261834

**Authors:** Leland B. Hyman, Clare R. Christopher, Philip A. Romero

## Abstract

Experimental methods that capture the individual properties of single cells are revealing the key role of cell-to-cell variability in countless biological processes. These single-cell methods are becoming increasingly important across the life sciences in fields such as immunology, regenerative medicine, and cancer biology. Existing single-cell analysis methods are often limited by their low analysis throughput, their inability to profile high-dimensional phenotypes, and complicated experimental workflows with slow turnaround times. In this work, we present Single-cell Nucleic Acid Profiling in Droplets (SNAPD) to analyze the transcriptional states of hundreds of thousands of single mammalian cells. Individual cells are encapsulated in aqueous droplets on a microfluidic chip and the content of each cell is profiled by amplifying a targeted panel of transcriptional markers. Molecular logic circuits then integrate this multi-dimensional information to categorize cells based on their transcriptional profile and produce a detectable fluorescence output. SNAPD analyzes over 100,000 cells per hour and can be used to quantify distinct cell types within populations, detect rare cells at frequencies down to 0.1%, and enrich specific cell types using microfluidic sorting. SNAPD provides a simple, rapid, low cost, and scalable approach to study complex phenotypes in heterogeneous cell populations.

## Introduction

The complex phenotypes displayed by multicellular organisms arise from interactions between highly specialized cell subpopulations. This intercellular heterogeneity plays a pivotal role in physiological, developmental, and disease processes. Transcriptomic analyses have traditionally been performed on the bulk collection of cells within a biological sample, and thus report the average transcriptional state of the population. This average does not capture the individuality of single cells within the population^1^. Single-cell transcriptomic approaches such as single-cell RNA sequencing (scRNA-seq) are helping to elucidate the role of cellular heterogeneity in fields ranging from immunology to oncology. These methods have revealed the continuum of individual cells states during differentiation processes such as hematopoiesis and secondary tumor formation^2,3^. They have also been used to characterize cellular heterogeneity in tumor microenvironments, where small subpopulations of cancer stem cells can drive disease progression and resistance to treatment^4^. Single-cell approaches are essential for studying biological processes that involve heterogeneous cell populations.

Existing single-cell transcriptional profiling methods often have low throughput, can only analyze a limited number of markers, and have laborious and expensive workflows. A widely used tool is scRNA-seq, where individual cells are isolated and subjected to next-generation sequencing. While this method theoretically allows one to measure any transcript in many individual cells, there is a tradeoff between transcript sequencing coverage and overall cell throughput. Consequently, scRNA-seq typically analyzes fewer than 10,000 cells per experiment^5^. Some applications may require analyzing a larger number of cells at the cost of profiling fewer transcriptional markers. Screening methods such as flow cytometry-based fluorescent in situ hybridization (Flow-FISH^6^) and microfluidic PCR-activated cell sorting (PACS^7,8^) can analyze a few transcripts in millions of single cells. However, these methods require laborious and involved sample processing that can span over multiple days. Thus, despite these recent technological advances, there is still need for new single cell transcriptional profiling methods that are low cost, simple/rapid, and can be scaled to millions of cells and hundreds of transcripts.

In this work, we demonstrate a highly scalable and streamlined technique for profiling the transcriptional states of single mammalian cells. We take advantage of recent advances in droplet microfluidics and molecular detection to develop the novel SNAPD (Single cell Nucleic Acid Profiling in Droplets) platform that is capable of profiling several mRNAs simultaneously across hundreds of thousands of mammalian cells. Our method starts by encapsulating single cells into microdroplets, followed by isothermal amplification of target RNAs to produce a fluorescent signal. The resulting droplets can be analyzed directly to determine the proportion of a specific cell type within a heterogeneous cell mixture, or the droplets can be sorted to enrich a particular cell type and collect its DNA and RNA. Hundreds of droplets can be analyzed per second, allowing ultra-high-throughput analysis and sorting. SNAPD can be multiplexed to examine multiple RNAs simultaneously, and these multidimensional signals can be integrated via molecular computation to capture more complex cellular phenotypes. The SNAPD workflow is highly streamlined—live cells are loaded directly into a microfluidic device with minimal processing beforehand and the entire transcriptional profiling process can be completed within a few hours. Due to its throughput, ease, and low cost, SNAPD provides a highly scalable and facile solution for single-cell transcriptional profiling.

## Results

### A microfluidic platform for transcriptional profiling of single cells

Droplet-based microfluidics is a powerful means to analyze single cells in high throughput. These systems encapsulate individual cells into picoliter-scale aqueous droplets, and each of these cells can be studied and analyzed in isolation from the bulk population. These techniques can achieve similar throughput to flow cytometry, but provide a more general format that can be adapted to a diverse array of biomolecular assays and readouts. For instance, droplet-based microfluidics has been used to perform single-cell RT-PCR to detect a specific mRNA transcript in tens of thousands of cells^7^. While this approach is powerful, it suffers from a fundamental problem with droplet-based assays: cells encapsulated and lysed in droplets will often result in high lysate concentrations that inhibit downstream assays such as PCR^9^. As a result, droplet-based PCR methods require complicated, multi-step workflows to remove reaction inhibitors, making assays difficult and less reliable. We therefore sought a simple droplet microfluidic method to analyze mRNA expression in hundreds of thousands of cells while avoiding PCR-based signal amplification altogether.

An ideal RNA amplification method for droplets could be performed in highly concentrated lysate with comparable sensitivity and selectivity to PCR. We identified Reverse Transcription – Loop Mediated Isothermal Amplification (RT-LAMP) as a method which satisfies these criteria. LAMP is an isothermal nucleic acid amplification method that uses a strand displacing polymerase (*Bst* Polymerase) to generate dumbbell DNA structures and exponentially extend them to longer concatemers^10^. Unlike DNA polymerases used in other methods, we found that *Bst* polymerase is resistant to lysate concentrations exceeding 10^6^ cells/mL (Fig. S1). LAMP-based assays have achieved limits of detection as low as 0.4 aM, which is comparable to nested PCR^11,12^. Additionally, RT-LAMP is isothermal, allowing for simple workflows that do not require thermocycling^13^. All of these favorable properties facilitated an unprecedentedly simple droplet-based LAMP workflow.

We developed SNAPD (Single cell Nucleic Acid Profiling in Droplets) to perform LAMP-based detection of specific mRNA transcripts from single cells (Fig. 1a). The SNAPD workflow is simple, rapid, and requires minimal hands-on processing. Live cells are stained with a live/dead indicator dye and loaded directly into a microfluidic device. Single cells are encapsulated into droplets containing LAMP and lysis reagents, along with target-specific primer sets. Droplets are then collected and incubated at a constant temperature, allowing the LAMP reaction to proceed. Finally, the droplets are reinjected onto a second microfluidic device, where their fluorescence is analyzed in high-throughput to detect LAMP-based amplification of the target markers.

**Figure 1.**
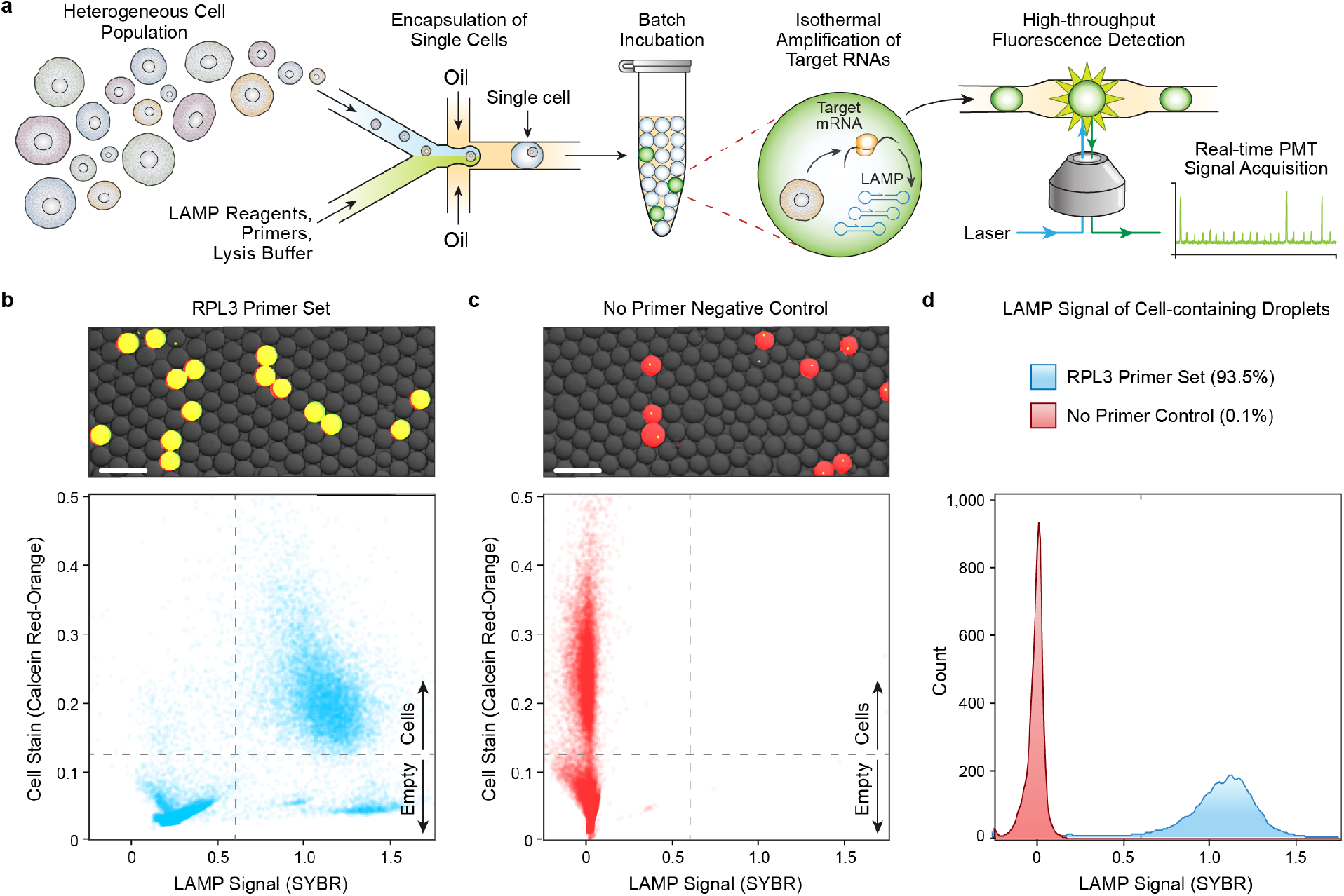
A microfluidic platform for high-throughput single-cell RNA profiling. (**a**) Schematic of the SNAPD workflow. Single cells are encapsulated into microdroplets with assay reagents, collected and incubated offline, and the fluorescence of each droplet is subsequently measured to indicate amplification of target RNAs. SNAPD can analyze the transcriptional state of over 100,000 cells per hour. (**b**) Microscopy image and scatterplot of SNAPD droplet fluorescence. We assayed MOLT-4 cells for expression of the housekeeping gene RPL3. Red fluorescence indicates the presence of a cell, while the green dye indicates target amplification. Droplets that appear yellow in the microscopy image contain cells and also displayed RPL3 amplification. Scale bars indicate a length of 200 μm. (**c**) Microscopy images and scatterplot of a SNAPD negative control with RPL3 primers omitted. Scale bars indicate a length of 200 μm. (**d**) Histograms of the cell-containing droplets SYBR fluorescence. Reactions containing the RPL3 primer set displayed 93.5% amplification, while the no primer negative control only amplified at 0.1%. These results demonstrate that SNAPD can reliably detect specific mRNA targets in single cells with high specificity.

We first benchmarked our SNAPD platform by measuring single-cell expression of the 60S Ribosomal Protein L3 (RPL3) housekeeping transcript. RPL3 is known to be uniformly expressed across single bone marrow progenitor cells^14^, and thus, we used the leukocytic leukemia line MOLT-4 as a positive control. We performed SNAPD on MOLT-4 cells with an RPL3 primer set, and also performed a no-primer negative control (Figs. 1b,c). Microscopy showed that cells (red) were loaded into approximately 10% of droplets, as expected, and nearly all cell-containing droplets displayed RPL3 amplification as visualized by SYBR green. In contrast, the no-primer negative control experiment displayed no SYBR green signal, indicating that LAMP-based mRNA amplification was responsible for the SYBR green signal in the cell-containing droplets.

We reinjected the SNAPD droplets onto a high-throughput fluorescence detection device, and measured the fluorescence of 10,000 cells. A majority of drops displayed a strong correspondence between the cell stain and LAMP amplification. There were a small fraction of drops that displayed LAMP amplification in the absence of cells, presumably due to free transcripts or dead cells that weren’t stained. There were also a small fraction of drops that displayed stained cells with no LAMP amplification. We gated the cell-containing droplets, and found that 93.5% of MOLT-4 cells amplified with the RPL3 primer set, as compared to 0.1% for the no-primer negative control (Fig. 1d). These results demonstrate that SNAPD can reliably detect specific mRNA targets in single cells. We were able to perform droplet generation and analysis at a rate of 300 Hz, allowing SNAPD to transcriptionally characterize over 100,000 cells per hour.

### Quantifying single-cell gene expression in heterogeneous cell populations

Single cell analysis methods can be used to distinguish different cell types and characterize heterogeneous cell populations. We used SNAPD to evaluate the expression of the HER2 (*ERBB2*) breast cancer marker in two cell lines. The SK-BR-3 breast cancer line overexpresses this gene, while the MOLT-4 leukemia line has no detectable expression^15^. We performed SNAPD on pure cell lines and found 97.1% of the SK-BR-3 cells displayed *ERBB2* expression, as opposed to 0.1% of the MOLT-4 cells (Figs. 2a,b). We also tested whether SNAPD could distinguish different cell types from the same tissue of origin by evaluating estrogen receptor (*ESR1*) expression in SK-BR-3 and MCF7 breast cancer lines. We found *ESR1* was expressed in 1.9% of SK-BR-3 and 67.9% of MCF7 cells (Fig. S2). *ESR1* is known to be heterogeneously expressed across single cells^16^.

**Figure 2.**
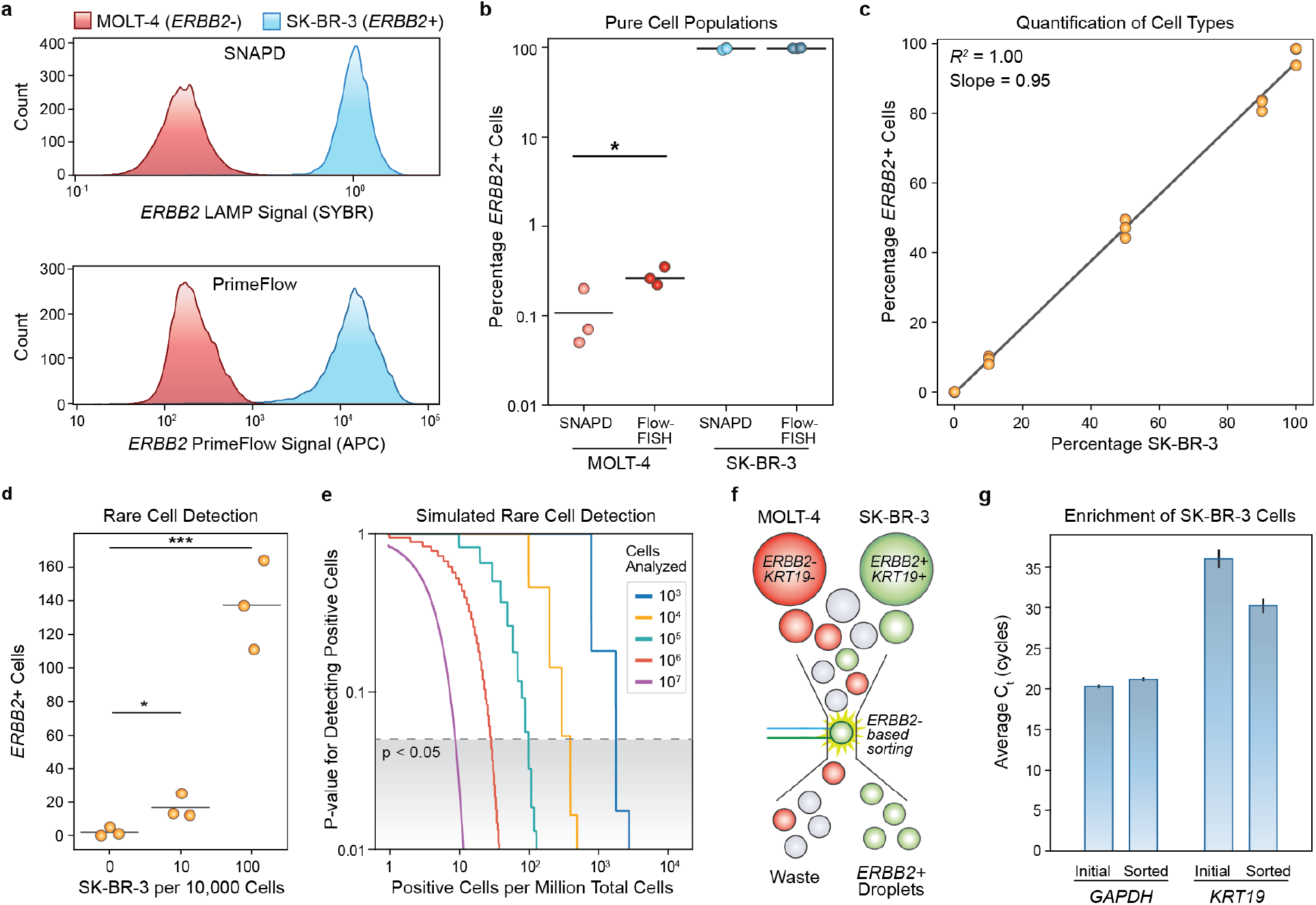
Quantification and enrichment of specific cell types. (**a**) SNAPD and PrimeFlow histograms comparing *ERBB2* (HER2) expression in MOLT-4 and SK-BR-3 cells. The overall fluorescence distributions are nearly identical, indicating SNAPD is comparable with well-established and commercially available single-cell analysis methods. (**b**) The percentage of *ERBB2+* cells as measured by SNAPD and PrimeFlow. Experiments were performed in triplicate. There was no significant difference between SNAPD and PrimeFlow when analyzing SK-BR-3 cells, while SNAPD displayed a lower false positive rate when analyzing MOLT-4 cells (p < 0.05). (**c**) SNAPD quantification of *ERBB2+* cells in mixtures of MOLT-4 and SK-BR-3 cells at varying proportions. Each measurement was performed in triplicate. These results indicate that SNAPD can quantify specific cell types across a broad range of proportions with high accuracy and precision. (**d**) SNAPD can detect rare *ERBB2+* cells in mixtures containing low proportions of SK-BR-3 cells spiked into MOLT-4 cells. All experiments were performed in triplicate. A mixture containing 0.1% SK-BR-3 cells was distinguishable from a sample of pure MOLT-4 cells (p < 0.05), indicating a limit of detection below 1 in 1,000 cells. (**e**) Simulation results exploring how the total number of cells analyzed affects SNAPD’s ability to detect rare cells. Analyzing 100,000 cells should be sufficient to detect positive cells at a prevalence of 1 in 10,000. (**f**) SNAPD can enrich specific cell types from mixtures using microfluidic sorting. MOLT-4 and SK-BR-3 cells were mixed at a 9:1 ratio, analyzed for ERBB2 expression using SNAPD, and droplets displaying *ERBB2* amplification were isolated using a microfluidic droplet sorting device. (**g**) SNAPD-based cell enrichment of SK-BR-3 cells was validated by performing RT-qPCR on the SK-BR-3-specific *KRT19* marker. The initial and sorted populations displayed similar *GAPDH* levels, indicating similar cell loading, while the sorted sample displayed a lower CT value for *KRT19*, indicating a higher proportion of SK-BR-3 cells. qPCR measurements were performed in triplicate and the error bars represent one standard deviation from the mean.

To further validate these results, we benchmarked SNAPD against Thermo Fisher’s PrimeFlow™ RNA assay kit, a well-established and commercially available Flow-FISH method to measure gene expression in single cells. PrimeFlow detects RNA by fixing cells, hybridizing fluorescent probes, washing away excess probes, and analyzing the target RNA within each cell by flow cytometery. SNAPD and PrimeFlow displayed highly similar *ERBB2* gene expression profiles across the two cell lines (Figs. 2a,b). There was no statistically significant difference between the SK-BR-3 (*ERBB2+*) detection rates of the two methods, although SNAPD did show significantly lower amplification for the MOLT-4 (*ERBB2-*) cell line (p<0.05). This implies that SNAPD can reliably identify *ERBB2* positive cells, and it may also have a lower false positive rate than PrimeFlow. We observed a small number of false positives in negative controls lacking *ERBB2* primers, which we believe are caused by cellular autofluorescence or events where the laser directly strikes a cell’s nucleus (Fig. S3).

After verifying that SNAPD could distinguish different cell types, we evaluated its ability to quantify specific cell types within a heterogeneous cell population. To do this, we combined SK-BR-3 and MOLT-4 in varying proportions, and used SNAPD to analyze the *ERBB2* marker in these defined cell mixtures. We found that SNAPD could reliably quantify the percentage of SK-BR-3 cells with a near-perfect linear fit (R^2^ = 1.00) and a slope of 0.95 (Fig. 2c). SNAPD provides a highly quantitative readout of a population’s cell types based on gene expression.

Single-cell methods can provide valuable information about rare cell types within a population. We tested SNAPD’s ability to detect rare cells by spiking a small known quantity of SK-BR-3 cells into MOLT-4 suspensions, and counting the resulting number of *ERBB2+* cells within the sample. SNAPD could reliably count SK-BR-3 cells in mixtures containing as few as 10 SK-BR-3 cells per 10,000 total cells analyzed, and could distinguish these mixtures from a negative control containing 100% MOLT-4 cells (Fig. 2d). SK-BR-3 cells at a prevalence of 1 in 10,000 total cells analyzed could not be distinguished from the negative control (Fig. S4). In this case, the total number of cells analyzed (10,000) limits SNAPD’s ability to detect rare cells. We built a simple model based on SNAPD’s cell classification parameters to explore how the total number of cells analyzed affects rare cell detection (Fig. 2e). We found that rare cell detection is highly dependent on screening throughput. We estimate our *ERBB2* SNAPD assay should be able to detect positive cells at a prevalence of 1 in 10,000 by screening only 100,000 cells total. Based on these results, SNAPD is capable of enumerating specific cell types from mixtures based on their RNA content, and has sufficient sensitivity and selectivity for a myriad of rare cell detection applications.

Many applications require isolating a specific cell type from a population; this enriched sample can then be analyzed using other methods such as mass-spectrometry or RNA-seq. The SNAPD single-cell assay can be combined with high-throughput droplet sorting to isolate specific cell types based on their gene expression profile. With a droplet microfluidic sorting device adapted from a previous design (Fig. S5)^17^, we used the *ERBB2* marker to enrich SK-BR-3 cells from an initial population containing 90% MOLT-4 and 10% SK-BR-3 (Fig. 2f). We then performed RT-qPCR on the initial and sorted pools to verify enrichment of SK-BR-3 cells. We used *GAPDH* as a reference gene, and estimated the samples’ proportion of SK-BR-3 cells by measuring the relative expression of the SK-BR-3-specific marker *KRT19*. We found the *GAPDH* levels were similar between samples, while the sorted sample contained higher *KRT19* levels, indicating a higher proportion of SK-BR-3 cells (Fig. 2g). Crude cell lysate in qPCR reactions resulted in poor amplification efficiency, and therefore we could not accurately quantify enrichment values. We estimated enrichment by taking each target’s observed qPCR efficiency into account following established qPCR calculations^18^ and by assuming GAPDH expression in SK-BR-3 cells is greater than or equal to its expression in MOLT-4 cells^15^. With these assumptions, we estimate that microfluidic sorting achieved at least a 3.2-fold enrichment of SK-BR-3 cells over MOLT-4 cells. These results demonstrate that SNAPD can be combined with microfluidic droplet sorting to isolate cell types from a heterogeneous population, and the RNA from the enriched cells can be recovered for downstream applications.

### Multiplex SNAPD to profile multiple RNA targets

Analyzing multiple RNA targets in single cells would allow SNAPD to identify more complex cellular phenotypes and classify cells with greater selectivity and sensitivity. Multiplex SNAPD requires simultaneous LAMP-based amplification of multiple RNA targets, in addition to orthogonal fluorescent readouts of each amplified target. Coupling LAMP amplification to a specific fluorescence signal is challenging due to the unpredictable and heterogeneous mix of concatemers generated by LAMP^10^.

We designed a polymerase-driven DNA strand displacement scheme to detect LAMP products in a sequence-specific manner (Fig. 3a). The goal was to build a molecular transducer that would convert a LAMP dumbbell product into a short DNA oligo that could then activate a fluorogenic reporter, or be fed into additional downstream strand displacement reactions. The molecular transducer consists of a ‘gate’ strand hybridized with an ‘output’ strand to produce a DNA duplex with a 3’ overhanging ‘toehold’. The LAMP dumbbell can hybridize to the transducer’s toehold, and its 3’ end acts as a primer for the polymerase to replicate the gate strand. The polymerase displaces the output strand during extension, releasing it. The free output strand can then activate a quenched fluorogenic reporter by displacing the quencher strand from the fluorophore strand. By designing a unique transducer for each LAMP product and a unique reporter for each transducer, we can link each RNA target’s amplification to a unique fluorophore. This detection strategy allows flexible multiplexing with minimal crosstalk.

**Figure 3.**
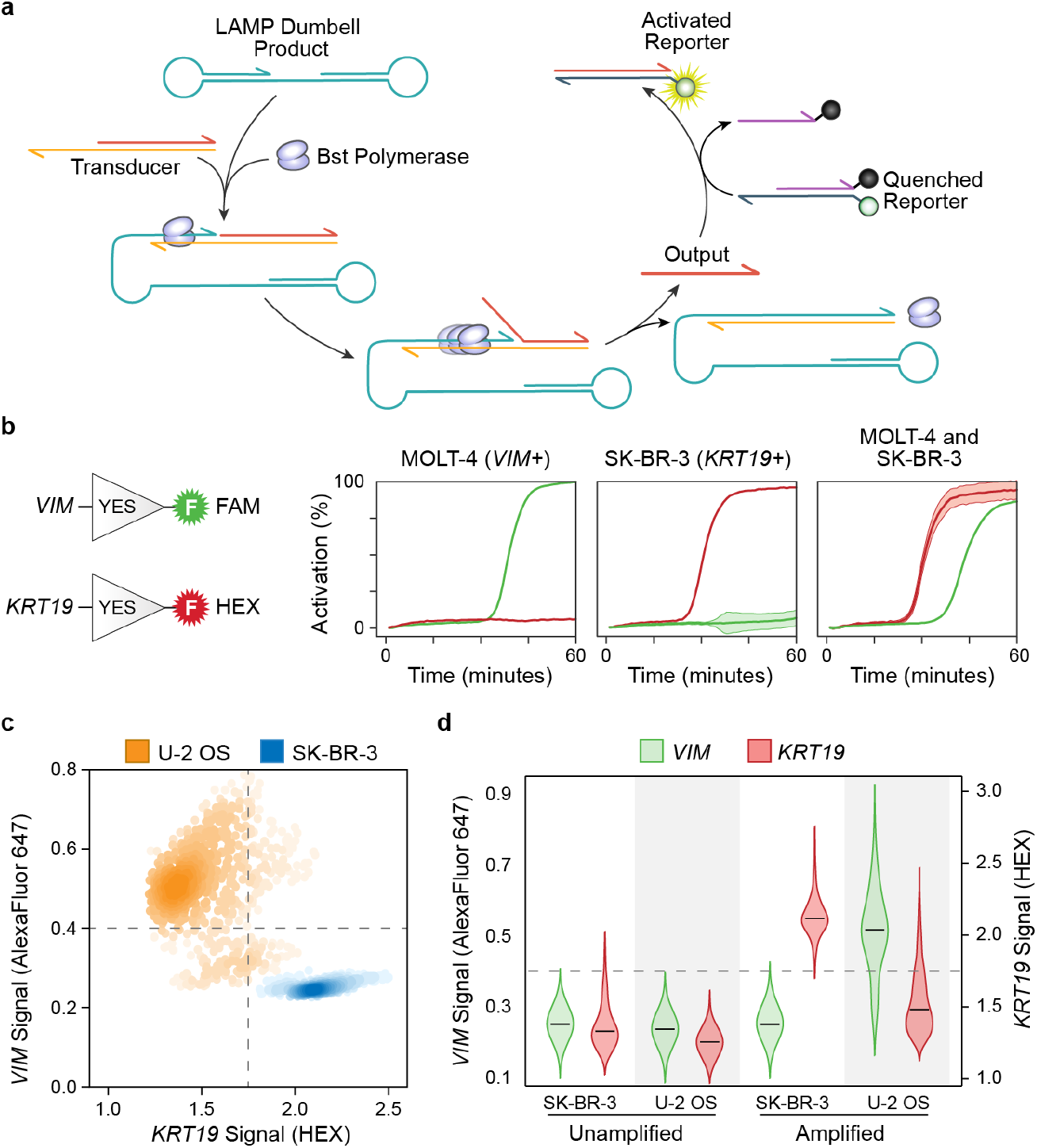
Multiplexed SNAPD profiles multiple mRNA targets in single cells. (**a**) Molecular transducer that converts the LAMP dumbbell product to a fluorescent readout in a sequence-specific manner. The *Bst* polymerase that is present in the LAMP reaction drives the displacement of the output strand; this output strand can then activate a fluorogenic reporter through toehold-mediated strand displacement. (**b**) Sequence-specific molecular transducers can operate orthogonally to link amplification of specific RNA targets to unique fluorescent outputs. We designed transducers that produce FAM fluorescence in response to *VIM* amplification and HEX fluorescence in response to *KRT19*. We added LAMP primer sets targeting *VIM* and *KRT19* and the two molecular transducers to lysates from either MOLT-4 or SK-BR-3 cells. The fluorescence of each reaction was monitored over time. The MOLT-4 lysate displayed specific activation of the VIM channel, while the SK-BR-3 lysate activated the KRT19 channel. A mixture of the two lysates activated both channels. We performed each measurement in triplicate. (**c**) Simultaneous profiling of *VIM* and *KRT19* expression in single cells using SNAPD. We combined the orthogonal *VIM/KRT19* transducers and SNAPD to analyze SK-BR-3 cells and the mesenchymal cell line U-2 OS. 98.4% of SK-BR-3 cells occupy the *VIM-/KRT19+* quadrant, while 83.7% of U-2 OS cells fall into the *VIM+/KRT19*-quadrant. (**d**) Violin plots showing the marginal distributions of VIM and KRT19 signal. SK-BR-3 cells display low *VIM* and high *KRT19* expression, while U-2 OS cells display high *VIM* and low *KRT19*. Also shown are the droplet fluorescence distributions before amplification to demonstrate the observed signals are a result of LAMP.

We tested our LAMP detection scheme and its ability to be multiplexed by designing two orthogonal amplification/transducer/reporter systems. The first produces a FAM signal in response to the mesenchymal marker vimentin (*VIM*), and the other produces a HEX signal in response to the epithelial marker cytokeratin 19 (*KRT19*). We added these orthogonal transducers to lysate from MOLT-4 (*VIM+/KRT19-*) and SK-BR-3 (*VIM-/KRT19+*) cells. As designed, the MOLT-4 lysate produced a FAM signal only, the SK-BR-3 produced a HEX signal only, and a 1:1 mixture of the lysates activated both channels (Fig. 3b). These results demonstrate that LAMP can amplify multiple targets in highly concentrated cell lysate, and that our transducer designs operate orthogonally to activate two separate fluorophores without signal crosstalk.

We combined our orthogonal LAMP transducers and SNAPD to evaluate the expression of *KRT19* and *VIM* in single cells. These two markers indicate a cell’s position along the epithelial-mesenchymal axis, and thus provide valuable information about cellular differentiation and cancer invasiveness^19^. We analyzed the epithelial-derived SK-BR-3 (*VIM-/KRT19+*) cells and found that 98.4% fell into the *VIM-/KRT19+* quadrant (Figs. 3c,d). We also performed an unamplified negative control to identify the signal background. Next, we analyzed the mesenchymal cell line U-2 OS (*VIM+/KRT19-*) and found that 83.7% fell into the *VIM+/KRT19*-quadrant. These results demonstrate that SNAPD can be multiplexed and used to profile multi-gene phenotypes in single cells.

### Analysis of Complex Phenotypes with Molecular Logic

Cellular phenotypes often depend on expression profiles across a multitude of genes. Our transducer-based multiplexing strategy is highly scalable; however, crowding within the optical spectrum restricts the number of orthogonal fluorescent reporters. To overcome these limitations, we devised a molecular logic scheme to integrate signals from multiple LAMP reactions and return a single fluorescent output based on a multi-level logical computation.

We used the *Bst* polymerase-driven strand displacement mechanism to design molecular logic gates comprising YES, NOT, OR, AND, and AND-NOT operations (Figs. 4a-e, S6). These designs build off of the LAMP transducer mechanism described in the previous section (Fig. 3a). The YES gate is simply a transducer and converts a LAMP product to a fluorescent output. The NOT gate inverts a LAMP signal by passing the output of a transducer to displace an unquenched fluorophore, which can then hybridize to a strand containing a quencher. The OR gate is composed of two transducers which accept different input sequences, but produce identical output strands. The presence of either LAMP signal will produce the output and generate a fluorescent signal. The AND gate similarly incorporates two transducers, but also contains a ‘threshold strand’ which sequesters the transducers’ output. When only one of the transducers is activated, the threshold strand sequesters all of the output; however, activation of both transducers produces a stoichiometric excess of the output strand over the threshold strand, leaving a sufficient amount of output strand to activate the fluorogenic reporter. Thus, a fluorescent signal is only produced when both LAMP signals are present. The AND-NOT gate is composed of a transducer that generates a fluorescent reporter in response to the first LAMP input, and is then followed by a not gate that inverts this signal in response to the second LAMP input. This gate will produce a fluorescent signal if the first input is on and the second input is off. This set of molecular logic gates is functionally complete, so in principle individual gates can be combined to produce any conceivable logical operation.

**Figure 4.**
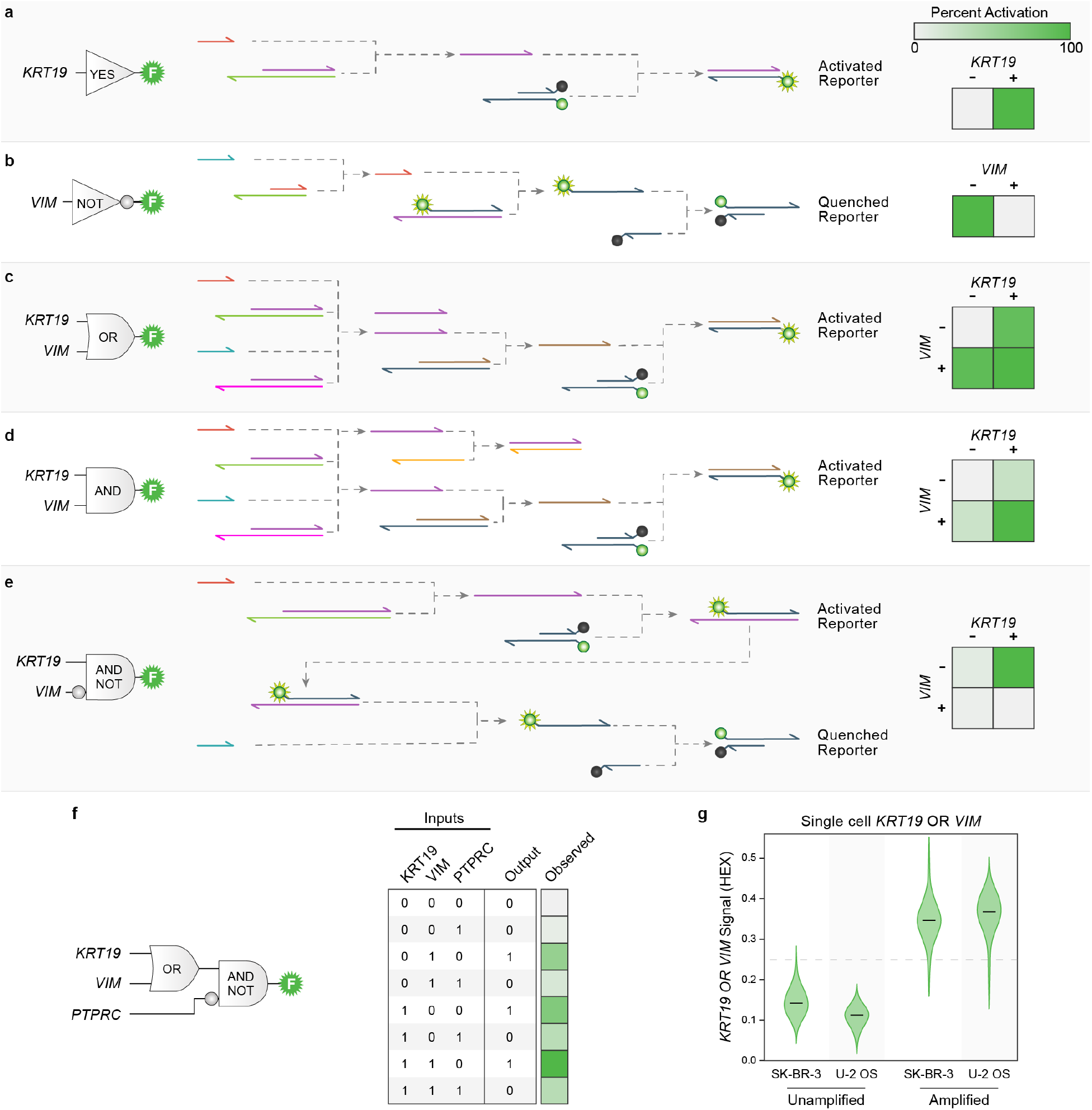
Molecular logic gates integrate multiple LAMP signals. (**a-e**) Molecular logic gate designs for YES, NOT, OR, AND, and AND-NOT operations. The designs leverage *Bst* polymerase’s strand displacement activity to drive the reaction cascade. We tested the designs on various combinations of purified RNA targets and all designs displayed the correct truth tables with a high signal-to-background ratio. (**f**) Multi-level logic circuit that integrates LAMP signals from *KRT19, PTPRC*, and *VIM* targets. The circuit displayed the intended logical outputs based on all 2^3^ combinations of inputs. (**g**) Integrating information across multiple markers in single cells. We applied a molecular OR gate to combine *KRT19* and *VIM* signals from single SK-BR-3 or U-2 OS cells. As expected, both cells activated the molecular OR gate, while an unamplified negative control displayed low fluorescence levels.

We constructed molecular logic gates to detect the presence of *VIM* and *KRT19* signals, and tested their ability to produce the expected truth tables. We added the molecular logic gates to LAMP reactions containing various combinations of purified RNA targets, and found the gates produced the desired logic with a high signal-to-background ratio. Across all gates, the average ON signal was 1.3-fold higher than the average OFF signal. The AND gate showed the lowest signal-to-background ratio with a 1.1-fold difference between the average ON state and the average OFF state.

We next built a multilevel logic cascade to integrate three LAMP inputs and activate a single fluorescent reporter (Figs. 4f, S6). We designed this circuit to implement (*KRT19* OR *VIM*) AND-NOT *PTPRC* (CD45), a logical operation which could be used to distinguish epithelial or mesenchymal circulating tumor cells from leukocytes in blood samples^20^. We found that this circuit correctly identified each combination of RNA inputs with a signal-to-background ratio of (1.2). This demonstrates that our molecular logic gates can be stacked for profiling higher-dimensional cell phenotypes.

We combined the molecular OR gate and SNAPD to evaluate whether single cells express *KRT19* or *VIM* (Fig. 4g). As expected, both SK-BR-3 cells (*KRT19+/VIM-*) and U-2 OS cells (*KRT19-/VIM+*) produced a fluorescent output. This shows that our LAMP based logic gates can analyze single cells in droplet reactions. Together, these logic gates provide a powerful means to enhance SNAPD and enable multiple transcripts to be profiled with a single fluorescence output.

## Discussion

Multicellular organisms are composed of hundreds of distinct cell types that interact to drive physiological, developmental, and disease processes. Single-cell analysis methods provide a valuable means to study specific cell types within heterogeneous cell populations. Single-cell approaches have been applied to a myriad of biological topics: dissecting heterogeneity in solid tumors^21^, elucidating drivers of cell fate during embryogenesis^22^, and reconstructing transcriptional regulatory networks^23^, among others. In this work, we developed Single-cell Nucleic Acid Profiling in Droplets (SNAPD) to analyze the transcriptional states of hundreds of thousands of single mammalian cells. SNAPD provides a simple, low cost, rapid, and scalable approach to study complex phenotypes in heterogeneous cell populations.

Existing single-cell transcriptomic methods include single-cell RNA Sequencing (scRNA-seq), single-cell qPCR (sc-qPCR), *in situ* hybridization methods such as Flow-FISH, and PCR-Activated Cell Sorting (PACS) in droplets. While methods such as scRNA-seq and sc-qPCR can analyze thousands of different transcripts, they suffer from low throughput (typically <10,000 cells)^24^, complicated and expensive workflows, and long turnaround times. On the other end of the spectrum, Flow-FISH and PACS profile smaller numbers of transcripts across millions of cells^8^. However, Flow-FISH and PACS both suffer from expensive and complex workflows, and are limited in their ability to analyze multiple transcripts. SNAPD occupies a unique position among single-cell analysis methods by providing high throughput with the potential to analyze multiple transcripts. SNAPD can analyze over 100,000 cells per hour, providing throughputs comparable to Flow-FISH and PACS. SNAPD’s analysis throughput could be reasonably scaled tenfold by optimizing the droplet size and device flow rates^17,25^. Furthermore, when combined with molecular logic, SNAPD can detect multiple transcripts and integrate these signals to produce a single fluorescence output. In principle, this molecular logic strategy could be scaled to process 10s of transcriptional inputs. Finally, SNAPD’s experimental workflow is highly streamlined, requires minimal hands-on processing, and provides results within a few hours.

We evaluated SNAPD’s ability to detect specific cell types with high sensitivity and specificity by analyzing defined mixtures of *ERBB2*-positive and -negative cells. We found SNAPD could reliably quantify the proportion of positive cells across a range spanning from 0.1% to 100%. Furthermore, we compared SNAPD to Thermo Fisher’s well-established Flow-FISH (PrimeFlow™) kit and found the two methods detected the same number of *ERBB2*-positive cells. However, SNAPD displayed nearly a 3-fold lower false positive rate for *ERBB2*-negative cells, suggesting that it has substantially higher specificity than Flow-FISH. This low false positive rate allowed us to detect rare cells from mixed populations with exquisite sensitivity and specificity. We successfully detected *ERBB2*-positive cells at 0.1% prevalence in a mixture, suggesting SNAPD’s limit of detection is less than 1 target cell per 1,000 total cells. Based on our results, we predict that *ERBB2*-positive cells could be reliably detected at a prevalence of 0.01% if the number of analyzed cells were increased from 10,000 to 100,000 (Fig. 2e). This level of sensitivity would be sufficient for numerous rare cell detection applications, including isolating mesenchymal stem cells from bone marrow^26^, pericytes from adipose tissue^27^, and breast cancer progenitors from pleural effusions^27^. These are therefore highly tractable future applications for our SNAPD platform.

In this work, we developed a novel multiplex LAMP detection scheme to analyze multiple RNA targets simultaneously. Previous efforts to perform sequence-specific detection of LAMP products have relied on a branch migration-based strand displacement mechanism targeting the loop region of the LAMP dumbbell structure^28,29^. Our LAMP detection scheme also uses a strand displacement mechanism, but takes advantage of the LAMP reaction’s *Bst* polymerase to drive strand displacement using the 3’ end of the LAMP product as a primer. The displaced strand can then displace a DNA duplex containing a fluorophore:quencher pair, thereby producing a fluorescent signal. Polymerase-driven strand displacement should display faster kinetics than branch migration because it does not rely on a random-walk strand exchange process^30,31^. In addition, the rate of polymerase-driven strand displacement depends only linearly on the length of the transducer’s duplex region, allowing longer duplex domains that result in less background leakage. Our polymerase-driven strand displacement scheme enables fast, robust, and orthogonal detection of specific LAMP products, even in complex mixtures. This capability was essential for our SNAPD platform to analyze the expression of multiple transcripts from single cells, but could also find more general applications in point-of-care nucleic acid detection.

We designed and implemented a novel set of DNA logic gates to integrate multiple LAMP signals into a single fluorescent output. Our designs leverage *Bst* polymerase activity to drive strand displacement, resulting in simple and robust DNA logic gates with fast kinetics. A similar DNA logic approach was recently described that used *Bst* polymerase to drive chemical reaction networks consisting of single-stranded oligos and primers^32^. Individual DNA logic gates can be combined into multi-input, multi-layer circuits that perform complex logical operations on nucleic acid inputs^32–35^. This would allow us to design logic circuits to identify very specific cell types based on the expression of multiple genes. For example, circulating tumor cells can be separated from background leukocytes and divided into specific epithelial or tumor generating subtypes based on the expression of 5 markers: EpCAM, CD44, CD34, CD45, and 7AAD^36^. This would preclude usage of other methods such as PrimeFlow™, which only analyzes up to 4 markers. DNA logic gates can also be used to build digital encoders that convert *2^n^* binary inputs to *n* distinct outputs^37^. We could apply digital encoders to profile gene expression states of 16 different transcripts using only four fluorescence channels. This would allow enumeration of large numbers of distinct cell phenotypes in heterogeneous populations, while maintaining a simple optical detection setup. The modularity, flexibility, and robustness of our polymerase-driven DNA logic gates will enable numerous applications of SNAPD.

Our SNAPD single-cell analysis platform has several limitations that could be addressed in future iterations. First, we found LAMP primer design to often require screening multiple designs to identify a primer set with fast amplification and low background. Some of this variability is likely caused by mRNA secondary structure or the specific splice isoforms present in each cell type. To overcome this, we found it was useful to include two redundant LAMP primer sets targeting each transcript. This strategy increased the overall LAMP signal and resulted in a lower false negative rate in SNAPD. To ensure successful amplification, we also targeted exonic regions shared between highly expressed splice variants. However, a systematic primer design framework that considers mRNA secondary structure across splice isoforms may improve the reliability of our method.

Another potential limitation involves measuring poorly expressed mRNA targets, an issue which is common to single-cell RNA sequencing assays^38^. Some markers may only have 1-10 transcripts per cell and result in low femtomolar concentrations when diluted into a ~500 pL droplet. LAMP assays have achieved sub-femtomolar and even attomolar detection of nucleic acids^39,40^. However, it’s unclear if this high level of sensitivity will translate to our droplet assay, which operates in dense cell lysate and has a high surface-to-volume ratio due to the microemulsion environment. Many of the transcriptional markers tested in this work are considered highly expressed, with normalized expression (NX) values ranging from 49.5-284.6^15^. The lowest expressed marker tested was *ESR1*, with an NX value of 24.3 in MCF7 cells. This is only 5% of the highest expressed gene in the MCF7 cell line^15^. Future work should explore SNAPD’s ability to measure lowly expressed transcripts, and compare it to the high dropout rates observed in single-cell RNA-seq.

A final limitation involves the detection of ultra-rare cells such as circulating tumor cells (CTCs). CTCs are typically found in blood samples at frequencies on the order of one CTC per 10 million nucleated blood cells^41,42^. We estimate that SNAPD is currently appropriate for enumerating rare cells at frequencies of one target cell per 1,000 total cells. To detect CTCs, we would need to reduce SNAPD’s false positive rate by approximately three orders of magnitude. We believe many false positive events arise when a cell nucleus passes through the laser, causing a large spike in the SYBR signal due to the genomic DNA. This could be addressed by using transducers instead of intercalating dyes, processing the time traces to filter large spikes, or examining the emission spectra in multiple channels to correct for cellular auto fluorescence^43^. Another source of false positive events came from amplification of extracellular RNA and/or DNA, which has been observed previously in droplet PCR^7,8^. We reduced the impact of these rogue nucleic acids by adding RNase I and DNase I to the input cell suspension, and then inactivated the nucleases with reducing agents during cell encapsulation into droplets. We also pre-treated the LAMP mastermix solution with DNase I to eliminate product contamination, and quenched with reducing agents prior to the assay. We found this approach reduced the false positive rate as much as 100-fold without reducing the true positive rate. Additional nucleases and other additives may decrease the false positive rate even further. Despite these various sources of false positive events, SNAPD remains a highly selective assay with an overall false positive rate that is significantly lower than established Flow-FISH methods.

Increasing SNAPD’s cell analysis throughput would improve rare cell detection and provide a more comprehensive view of heterogeneity within cell populations. The throughput of our system is currently limited by the microfluidic fluorimeter device, which processes droplets at a rate of approximately 300 Hz. Attempts to operate this device at faster flow rates resulted in droplet shredding due to the large size of the drops relative to the microfluidic channels. The system’s throughput could be increased by either redesigning the microfluidic device to better accommodate large droplets or by decreasing the droplet volume. We initially chose a relatively large 500 pL droplet size to mitigate LAMP reaction inhibition by concentrated cell lysate. However, lysate titration experiments (Fig. S1) suggest that we could decrease droplet sizes to at least 100 pL, which would increase the screening throughput five-fold. Decreasing the droplet size has the additional benefits of reducing reagent cost and analysis time, thereby further enabling larger samples and more accurate rare cell detection.

Experimental methods that capture properties of individual cells across large heterogeneous populations are essential for understanding the intricate and complex behaviors of biological systems. Single-cell analysis methods are currently hampered by complicated and arduous workflows, slow turnaround times, and expensive reagents. In this work, we developed the SNAPD platform to address the limitations of existing single-cell analysis methods. SNAPD is simple, rapid, low cost, and scalable, making it a powerful approach to characterize single-cell phenotypes. With these advantages, SNAPD could help lower the barrier to entry for single-cell methods and allow widespread adoption among life science researchers.

## Materials and Methods

### LAMP primer design

For each gene target, we first used data from the Human Protein Atlas Project^15^ to identify its most highly expressed transcriptional isoforms in our target cell type. We then used the Ensembl Genome Browser^44^ to identify common regions between highly expressed isoforms, and designed LAMP primers to target these sequence regions. We designed LAMP primers using PrimerExplorer V5 software^45^ (Eiken Chemical Co.). We found SNAPD’s sensitivity was greatly improved if we used two different sets of LAMP primers to target each target. In these cases, we designed primers such that little or no overlap occurred between them.

### DNA Complexes and Primer Mixes

We ordered all DNA oligos from Integrated DNA Technologies (Coralville, Iowa), and dissolved each into nuclease-free water (Thermo Fisher) prior to storage at −20 °C. On the day of an experiment, we prepared stocks of each DNA complex in DEPC-treated PBS by slowly annealing the strands from 97 °C to 23 °C at a rate of −2 °C/min. We then stored these DNA complexes on ice and protected from light until the time of experiment. We prepared LAMP primer mix stocks in nuclease-free water (Thermo Fisher) and stored at −20 °C.

### Production of mRNAs using *in vitro* transcription

DNA templates for CK19, VIM, and ERBB2 transcripts were synthesized by Integrated DNA Technologies (Coralville, Iowa). We cloned these genes into the pET-22b vector under a T7 promoter. We performed *in vitro* transcription using the HiScribe™ T7 High Yield RNA Synthesis Kit (New England Biolabs), and purified the resulting RNA using a GeneJET RNA Purification Kit (Thermo Scientific). We quantified each RNA sample’s concentration using a NanoDrop™ Spectrophotometer (Thermo Scientific) and stored RNA stocks at −80 °C in DEPC-treated PBS.

### Microfluidic device Fabrication

We fabricated microfluidic devices from polydimethylsiloxane (PDMS) via a soft photolithography process. We first deposited SU-8 3025 or 3010 photoresist onto a silicon wafer and spun to achieve the desired layer height. Next, we used a photomask to pattern microfluidic channels on the wafer and removed uncrosslinked regions with SU-8 developer. We then placed the patterned wafer into a petri dish, submerged in uncured PDMS (Dow Corning Sylgard^®^ 184) (11:1 polymer:cross-linker ratio), and cured at 72 °C for at least 1 hour. After PDMS polymerization, we excised the patterned PDMS device from the mold and bonded it to a glass microscope slide via plasma treatment. Finally, we treated the microfluidic channels with Aquapel (Pittsburgh Glass Works) to make the channels hydrophobic.

### Cell Culture and Staining

We subcultured MOLT-4 cells (American Type Culture Collection) in a 1:8 ratio every two days in RPMI-1640 Medium (Gibco) supplemented with 10% FBS (Gibco) and 1X Antibiotic-Antimycotic (Gibco). We subcultured SK-BR3 and U-2 OS cells (American Type Culture Collection) in a 1:4 ratio every two days and in DMEM, high glucose (Gibco) supplemented with 10% FBS and 1X Antibiotic-Antimycotic. On the day of experiment, we collected each cell type and washed twice with DPBS (Gibco). We then stained cells with 10 μM dye in DPBS for 30 minutes on ice. In droplet transducer and OR gate experiments, we stained with CellTrace™ Calcein AM (Invitrogen). In all other droplet experiments, we used CellTrace™ Calcein Red-Orange AM (Invitrogen). We then washed cells twice and resuspended in DMEM (Gibco) containing 12.4 U/uL RNase I_f_ (New England Biolabs) and 0.025 U/uL DNase I (Thermo Fisher). We performed the *ESR1* experiment (Fig. S2), multiplex transducer experiment (Fig 3cd), and droplet OR gate experiment (Fig. 4g) with an earlier protocol that did not include nuclease treatment and used DPBS instead of DMEM.

### Bulk LAMP and DNA logic assays

We performed bulk RT-LAMP and DNA logic assays on purified RNAs, cells, and mixtures of cells in triplicate using a Bio Rad CFX Connect qPCR machine. We incubated the reactions at 65 °C and monitored FAM, HEX, and/or SYBR fluorescence channels. Each reaction comprised a total volume of 10 μL, consisting of 1.6 μM each FIP/BIP primer, 0.2 μM each F3/B3 Primer, 0.4 μM each LoopF/B Primer, 1X WarmStart LAMP Master Mix (New England Biolabs), and 0.5 U/μL SUPERase•In™ RNase Inhibitor (Invitrogen). We added DNA complexes at varying concentrations given in Table S2. LAMP primer and logic gate sequences are shown in Table S1. In reactions without any DNA complexes, we included LAMP Fluorescent Dye (New England Biolabs) as a general LAMP indicator. For experiments performed on purified RNAs, we added *in vitro* transcribed *KRT19, VIM*, and/or *PTPRC* RNAs at the concentrations shown in table S2. For experiments performed on cells, we also included 2.5% Tween-20 (Sigma Aldrich) in the reaction buffer to act as a lysis reagent. We then added intact, unstained cells immediately before starting an experiment at a final concentration of 50 cells/μL per cell type.

### Single-cell experiments in microfluidic droplets

We used microfluidic dropmakers to encapsulate cells with LAMP components and lysis reagents into 500 pL droplets (Fig. S5). This dropmaker included 2-3 aqueous inlets depending on the experiment type. The device flow rates and inlet compositions are given in Table S3. Table S2 lists the final concentration of each logic gate for droplet logic experiments. We used standard LAMP primer concentrations, as described in ‘Bulk RT-LAMP Assays.’ In all experiments, we adjusted the cell concentration such that approximately one in every ten droplets contained a single cell. We collected droplets into a microcentrifuge tube and incubated at 65 °C for 1 hour prior to analysis. We then reinjected droplets onto a second microfluidic device with additional oil for spacing and measured their fluorescence in FAM/SYBR, HEX, and Alexa Fluor 647 channels. We performed subsequent gating and analysis in Flowjo software. For single-target experiments, we omitted all LAMP primers as a negative control. For multiplexed experiments, we performed unamplified negative controls that were only incubated at 65 °C for five minutes. We analyzed at least 10,000 cells for each replicate of each experiment, except for the droplet logic gate and *ESR1* experiments, where we analyzed over 1,000 cells.

During the course of SNAPD development, we made several refinements to reduce false positive events. We switched from a 3-inlet droplet generator to a 2-inlet design. We also added RNase If (New England Biolabs) and DNase I (Thermo Fisher) to the cell suspension to destroy extracellular DNA and RNA. Likewise, we pre-treated the LAMP mastermix with DNase I for 30 minutes at room temperature, followed by a 30 minute inactivation with DTT on ice, followed by primer addition.. We used the 3-inlet, no-nuclease method for *ESR1*, droplet transducer, and droplet OR experiments, and the 2-inlet method with nuclease treatment for all other droplet experiments.

### Droplet fluorescence data processing

Our microfluidic measurements produce fluorescence time traces of the droplets passing the laser. We used a fixed threshold to distinguish droplets from the carrier oil, and the average fluorescence within each droplet was recorded at multiple wavelengths. The resulting data looks similar to flow cytometry data with many events across multiple fluorescence channels. For single-target experiments, we performed signal compensation to reduce crosstalk between the FAM/SYBR and HEX channels. We then gated out excessively large/small drops by examining the droplet size (duration of signal) and removing the top and bottom 8% of events. Next, we selected the cell-containing droplet population by manually gating droplets displaying the calcein red-orange cell stain in the HEX channel. We then down-sampled 10,000 random events from the resulting droplet population to allow comparisons across samples. We defined a positive/negative LAMP amplification threshold as 2.8-fold higher than the median fluorescence of the empty droplet population. These same thresholds were applied to negative control conditions without LAMP primers. For the droplet logic experiment, we used a different optical configuration and different fluorophores, but used the same general compensation and gating strategy.

### RNA Flow-Fish

We performed assays in triplicate using the Primeflow RNA Assay Kit™ (Thermo Fisher) with Alexa Fluor 647 *ERBB2* target probes, following the recommended protocol. We stained MOLT-4 and SK-BR-3 cells with LIVE/DEAD™ Fixable Green Dead Cell Stain Kit, for 488 nm excitation (Invitrogen) prior to treatment. As a negative control, we also performed the entire protocol without the *ERBB2* target probes. We analyzed the samples on a BD Fortessa Flow Cytometer using APC and FITC channels, and performed subsequent gating and data analysis in FlowJo software.

### Simulating SNAP’s ability to detect rare cells

We used data from *ERBB2* limit of detection experiments (Fig. 2d) to estimate SNAPD’s false positive rate (FPR) as 0.02%. Similarly, We used data from SK-BR-3 *ERBB2* amplification (Fig. 2b) to estimate SNAPD’s true positive rate (TPR) as 97.1%. For each trace in Fig 2e, we fixed the number of cells analyzed, and simulated varying proportions of SK-BR-3 cells in a MOLT-4 background. In each simulation, we generated a binomial distribution to approximate the pure MOLT-4 population using the estimated FPR and the number of cells analyzed. We then calculated the expected number of observed positive cells to be equal to the number of MOLT-4 cells * FPR + the expected number of SK-BR-3 cells * TPR. We used this expected value to perform a binomial test against the pure MOLT-4 population and obtain a p-value for detecting positive cells.

### Enrichment of *ERBB2+* cells via microfluidic droplet sorting

We used droplet sorting to enrich *ERBB2+* cells from a cell mixture, and verified enrichment using qPCR. We performed *ERBB2* SNAPD on a 9:1 mixture of MOLT-4 and SK-BR-3 cells stained with CellTrace™ Calcein Red-Orange AM (Invitrogen). We then reinjected these droplets onto a dielectric sorting device (Fig. S5) with flow rates of 100 μL/hr for SNAPD droplets, 400 μL/hr for reinjection oil, and 1,000 μL/hr for bias oil. We measured the fluorescence of each drop in the FAM/SYBR and HEX channels. We sorted positive droplets containing cells by applying a series of 250 800-V, 10-kHz DC pulses across the sorting junction. We performed sorting experiments in triplicate.

To verify enrichment, we extracted RNA from the sorted droplets and performed RT-qPCR on *GAPDH* and *KRT19*. We froze the sorted droplet samples at −20 °C overnight to coalesce the emulsions and preserve RNA integrity. We then thawed samples, extracted the aqueous layer containing the RNA, and pooled samples from the three sorting replicates to increase the RNA yield. For the initial (unsorted) samples, we used a 150,000 cells/mL suspension containing a 9:1 mixture of MOLT-4:SK-BR-3 cells, and serially diluted this mixture in 2-fold intervals to identify an RNA concentration that matched the sorted droplets. We subjected this cell mixture to bulk *ERBB2* LAMP to match the experimental conditions of the sorted droplets. We then digested DNA in the sorted and initial (unsorted) samples by diluting 10-fold and adding 0.1 U/μL DNase I (Thermo Fisher) and 2 U/μL RNasin Plus RNase Inhibitor (Promega). We performed DNase digestion at 37 °C for 1 hour followed by a 95 °C inactivation for 5 minutes. We added 1 μL of the resulting DNase digestions to 10 μL RT-qPCR reactions (Luna One-Step Universal RT-qPCR Kit, New England Biolabs) containing 0.8 U/μL RNasin Plus (Promega) and *GAPDH* or *KRT19* primers (primer sequences listed in Table S1). For non-template controls, we added water instead of RNA. We performed thermocycling as recommended in the qPCR kit’s protocol. We performed all qPCR measurements in triplicate.

We calculated enrichment by evaluating how *KRT19* levels changed relative to the *GAPDH* reference gene. Since SK-BR-3 cells express the *KRT19* gene, a high *KRT19/GAPDH* ratio indicates SK-BR-3 enrichment relative to MOLT-4 cells. We derived an expression for enrichment based on a modified version of the standard 2^−ΔΔCT^ method. This expression accounts for varying PCR efficiencies between the target and reference gene^18^. Enrichment was calculated as

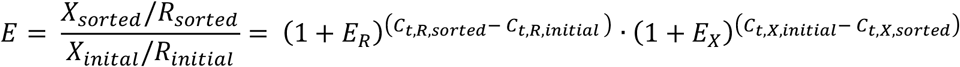

where *X*_sorted_ and *R*_sorted_ correspond to target (*KRT19*) and reference (*GAPDH*) RNA levels in the sorted sample, respectively; and *X*_inital_ and *R*_inital_ correspond to the initial sample. *C*_*t*,R,sorted_ is the number of PCR cycles for the reference gene to reach a defined threshold in the sorted sample, *C*_*t*,R,initial_ corresponds to *GAPDH* in the initial sample, *C*_*t,X*,sorted_ corresponds to *KRT19* in the sorted sample, and *C*_*t,X*,initial_ corresponds to *KRT19* in the initial sample. *E_R_* and *E_X_* are the PCR efficiencies for the reference and target, respectively; and were estimated to be *E_GAPDH_* = 0.36 and *E_KRT19_* = 0.17 by running qPCRs at varying cell lysate concentrations.

## Acknowledgements

P.A.R. is a Damon Runyon-Rachleff Innovator supported (in part) by the Damon Runyon Cancer Research Foundation (DRR-40-16).

## Author Contributions

L.B.H. and P.A.R conceived the project. L.B.H. and C.R.C. performed the experiments. L.B.H. analyzed the data with feedback from P.A.R.. L.B.H. and P.A.R. wrote the manuscript.

